# MetaNorm: Incorporating Meta-analytic Priors into Normalization of NanoString nCounter Data

**DOI:** 10.1101/2023.11.17.567577

**Authors:** Jackson Barth, Yuqiu Yang, Guanghua Xiao, Xinlei Wang

## Abstract

Non-informative or diffuse prior distributions are widely employed in Bayesian data analysis to maintain objectivity. However, when meaningful prior information exists and can be identified, using an informative prior distribution to accurately reflect current knowledge may lead to superior outcomes and great efficiency. We propose MetaNorm, a Bayesian algorithm for normalizing NanoString nCounter gene expression data. MetaNorm is based on RCRnorm, a powerful method designed under an integrated series of hierarchical models that allow various sources of error to be explained by different types of probes in the nCounter system. However, a lack of accurate prior information, weak computational efficiency, and instability of estimates that sometimes occur weakens the approach despite its impressive performance. MetaNorm employs priors carefully constructed from a rigorous meta-analysis to leverage information from large public data. Combined with additional algorithmic enhancements, MetaNorm improves RCRnorm by yielding more stable estimation of normalized values, better convergence diagnostics and superior computational efficiency. R Code for replicating the meta-analysis and the normalization function can be found at github.com/jbarth216/MetaNorm.

## 1. Introduction

### 1.1. Normalization methods for nCounter gene expression data

The medium-throughput platform NanoString nCounter has quickly become one of the most popular and efficient ways to analyze complex mRNA transcripts [8]. One reason for this is its ability to process formalin fixed, parrafin-embedded (FFPE) tissue samples effectively. While freshly frozen (FF) samples experience less degradation in storage, FFPE samples require less stringent storage requirements and are therefore cheaper and easier to retain [14]. This has led to a ubiquity of FFPE samples and has made the nCounter system an invaluable resource in medical research. Among other areas, the nCounter framework has been used to analyze FFPE samples in studies of colon, breast, and lung cancer [5, 10, 17]. However, a significant downside to using FFPE samples is the level of mRNA “modification” during the preservation and storage of the sample [11], causing higher levels of variability and uncertainty in the readings. Thus, it is critical that the gene read counts undergo an efficient normalization procedure before being formally analyzed.

In addition to degradation in FFPE samples, normalization can also help to remove background noise or lane-by-lane variation, which is common in FF samples as well. Perhaps the simplest approach to normalization is NanoStringNorm, an R package that implements NanoString guidelines and relies on summary statistics from the positive, negative and housekeeping probes to account for separate types of variation [16]. Other normalization methods, such as NAPPA^1^ and NanoStringDiff [18], use model-based approaches that can better account for the complexities in the data. Despite the improvements of NAPPA and NanoStringDiff, all three of these approaches use each type of probe in the same way, in that positive probes only remove lane-by-lane variation, negative probes only remove background noise, and housekeeping genes only remove variation in the amount of the input sample material. Among the latest normalization protocols to be developed is RCRnorm, which stands for **r**andom **c**oefficient hierarchical **r**egression **norm**alization [9]. RCRnorm is unique in that (1) it is an integrated system of hierarchical models for the different types of probes, (2) it is designed specifically to normalize FFPE data (but can be used with other types of samples), (3) it allows for sample-to-sample variation in the housekeeping genes [7], and (4) it uses a Bayesian framework for normalization, thus allowing statistical inference about key model parameters and uncertainty quantification for estimates while leading to better interpretability. Essentially, the RCRnorm model assumes that all probes in a given sample have a shared intercept and slope on a linear regression of log_10_ (gene read count) and log_10_ (RNA amount). While other probe-specific effects are included, this allows all types of probes to influence the normalization process, yielding more robust estimates (technical details of RCRnorm are outlined in section 1.3). In general, RCRnorm is much more flexible than its frequentist counterparts, with the Gibbs sampler able to explore the extremely high-dimensional parameter space and land on the targeted posterior distribution. The authors also show a significant improvement in the correlation between FFPE samples and their corresponding FF samples, which have gene read counts that are thought to be more accurate to the ground truth than the FFPE counts, with RCRnorm outperforming other normalization methodology at both the gene and patient level [9]. Additionally RCRnorm has also been found to remove more technical variation than its counterparts [3], including nSolver and NanoStringDiff.

### 1.2. Motivation to improve RCRnorm

Despite its impressive performance, RCRnorm is not without its drawbacks. The weakest point is its high computational cost, which is well-documented [3]. Large datasets require a significant amount of time and working memory to produce results, making RCRnorm an occasionally inconvenient choice for normalization. Another area of concern is the high number of parameters, which is typical in Bayesian hierarchical models, where it is common to rely on priors to regularize these parameters. Still, in a highly complex system, an abundance of parameters combined with an absence of accurate prior information can cause weak convergence. Further, a concern closer to the crux of the RCRnorm model is the construction of the prior distributions for the hyper-parameters *μ*_*a*_, *μ*_*b*_. These hyper-parameters represent the means of the sample-specific intercept and slope terms (respectively), which should reflect how the nCounter system inherently operates rather than the characteristics of an individual dataset. They have a significant impact on model estimates and so the results of RCRnorm are sensitive to the choice of prior distribution placed on these hyper-parameters. RCRnorm uses a jackknife approach on the positive probe data to estimate the prior mean and variance for *μ*_*a*_, *μ*_*b*_ (a normal distribution is assumed), so that the priors vary across datasets [9]. This makes the results of RCRnorm more sensitive to the size and quality of the dataset being normalized, since any data-based prior has no ability to mitigate potential bias that can arise from sparse or unreliable data. For these reasons, identifying and implementing new prior distributions for these quantities will improve model estimation, especially for messy or sparse datasets.

Since *μ*_*a*_, *μ*_*b*_ are important global parameters in the RCRnorm model, replacing the jackknife priors with non-informative priors would not effectively mitigate the potential bias and noise stemming from low quality data. Rather, *μ*_*a*_, *μ*_*b*_ reflects the underlying mechanism of the nCounter system and constructing an informative *a priori* distribution based on largely existing data of the same type will help avoid excessive data-dependence and provide external calibration to low quality data. In this big-data era, accessing data from multiple independent studies has become increasingly easy, due to federal regulations and substantial efforts made in data sharing. Thus, leveraging such information to construct informative yet objective priors becomes feasible.

There are two goals of this study: the first is to create these prior distributions via a rigorous meta-analysis of public FFPE gene expression datasets from independent studies using NanoString nCounter platform. To do so, we devise a Bayesian hierarchical model, which adopts the linear regression setup with random coefficients from RCRnorm. However, it is distinct from the RCRnorm model in several aspects. First of all, it deals with data from multiple studies and includes a layer to account for study-specific effects. Secondly, for reasons detailed in the beginning of section 2, this new model focuses on modeling data from positive control probes only. Thus, it is able to allow probe-specific variance terms, removing the simplifying assumption made by RCRnorm (i.e., constant variance across all positive controls). Thirdly, unlike RCRnorm, the new model, with much more data available in a meta-analysis, is able to account for the dependence between the sample-specific intercepts and slopes.

The second goal is to implement these informative prior distributions along with other important enhancements to improve the performance of RCRnorm. We refer to this new algorithm as MetaNorm. Among these enhancements are (1) optimized code and data structure, (2) implementation of constraints on the positive and negative probe residuals, and (3) a simplified sampling approach to normalized RNA expression estimates for housekeeping and regular genes. We show that these adjustments, along with the informative meta-analysis prior, improve computational cost, model convergence, prediction error and performance with low-quality data. In the remainder of this section, we outline in detail the original RCRnorm model [9]. In section 2, we provide an overview of our meta-analysis and a detailed summary of other improvements made to RCRnorm. In section 3, we compare the performance of MetaNorm with RCRnorm on four real-world datasets. We make a similar comparison to a traditional normalization algorithm in section 4 Finally, we provide a discussion of our findings in section 5.

### 1.3. Overview of the RCRnorm model

We conclude the Introduction with an overview of the notation used in this paper and a description of the RCRnorm model. Each FFPE dataset has *I* patient samples (indexed by *i*) with *P, N, H, R* positive probes, negative probes, housekeeping genes and regular genes (indexed by *p, n, h, r*), respectively. The datasets have a factorial structure, with each sample/gene combination represented exactly once per dataset (*I ·* (*P* + *N* + *H* + *R*) total observations). Typically, there are 6 positive probes (*P* = 6), a relatively small number of negative probes and housekeeping genes (i.e., *N <* 10 and *H <* 20) and a high number of regular genes (*R >* 80). Each of the six positive probes contains a known amount of mRNA, while the amount of mRNA in a regular or housekeeping gene is unknown and varies across samples. These mRNA levels must be estimated using the read counts. For all negative probes, target transcripts are absent and so their mRNA levels should be zero ideally. The aim of RCRnorm is to create consistent, unbiased estimates for this mRNA amount for every gene/sample combination that adjust for unwanted biological and technical effects. The remainder of this section summarizes the outline and technical details of the model [9].

The RCRnorm model is based on a series of integrated linear regression models with random coefficients. At the lowest level, the log_10_ gene read count *Y* for each of the four probe types is summarized in the below equations:

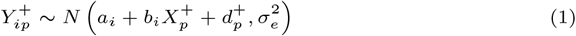

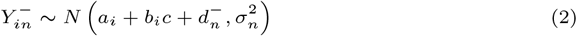

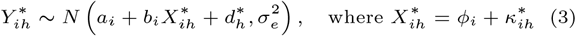

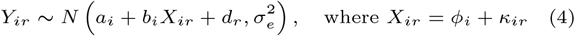

where the superscripts (+, −, *) indicate positive probes, negative probes, and housekeeping genes, respectively (no superscript in gene specific variables indicates regular genes); the log_10_ RNA levels are known and fixed for each positive probe 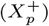 in all samples and are unknown but differ for all housekeeping and regular gene observations 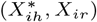; the unknown scalar *c* for negative probes can be interpreted as the mean non-specific binding level due to background noise. The intercept and slope terms (*a*_*i*_, *b*_*i*_) reflect how the log read count *Y* relates to the log RNA level *X* in each sample, and are assumed to be independent and identically distributed normal variables: 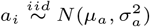 and 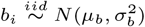. Note that for a given sample, these terms are shared by all the four categories and so occur in all four equations. Whereas *ϕ*_*i*_ represents the sample specific degradation level (with the constraint 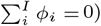, *κ*_*ir*_ and 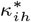 signify sample/gene specific mRNA levels before degradation (these parameters can be thought of as the model output), with 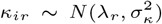 and 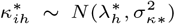. The variables 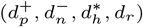 represent deviations from the general linear pattern and can be thought of as probe-specific residuals, with 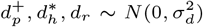 and 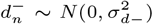. Jia et al. [9] suggests that all probe types have similar variability except for the negative probes, which tend to have higher variances, hence 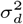 vs. 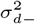 and similarly, 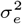 vs. 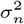 in (1)–(4). The left panel of figure 1 summarizes the hierarchical structure of the RCRnorm model. Note that the hyper-parameters *μ*_*a*_ and *μ*_*b*_ have data-based jackknife priors, which we seek to improve with the results of our meta-analysis. See sections 3 and 4 of Jia et al. [9] for more details of RCRnorm.

## 2. Methods

The first step in improving RCRnorm is to identify realistic, informative prior distributions to replace the current priors for *μ*_*a*_ and *μ*_*b*_. We maintain the assumption that the priors are independent normal distributions (the conjugate priors for *μ*_*a*_ and *μ*_*b*_), but rather than relying on the data that is currently being analyzed, the MetaNorm prior comes from a meta-analysis. We identified multiple independent NanoString nCounter gene expression datasets of FFPE samples from past studies and designed a Bayesian hierarchical model for meta-analysis to attain parameter estimates for the prior mean and variance of *μ*_*a*_ and *μ*_*b*_ irrespective of a specific dataset. For positive control probes, both *Y* (the log transformed count) and *X* (input RNA amount) are known so that we have strong information from the data about the intercept and slope terms. In contrast, for housekeeping and regular genes, only *Y* is known and *X* is latent with a complex underlying structure, so such information is much weaker. Since our main interest is on the intercept and slope terms, the meta-analysis only considers these positive controls (6 per sample) to find the posterior distributions for *μ*_*a*_ and *μ*_*b*_. As will be shown later, this would also allow us to remove simplifying assumptions made by RCRnorm for the purpose of estimation stability. The remainder of this section outlines the design, data and results of this meta-analysis.

### 2.1. Datasets

We identified 17 potential datasets containing mRNA FFPE samples from human subjects, 14 identified in the gene expression omnibus with search terms “mRNA”, “FFPE” and “nCounter” and 3 identified through collaborators or made available by authors on github. Details for each dataset can be found in table S1 of the supplementary material [1]. Two of these datasets (id 4 and 5) were used for testing the RCRnorm model, and were removed from the meta-analysis pool in an attempt to be completely independent from the original study. Two other datasets were removed for having low data quality, with one having a high number of low-quality samples by the NanoString quality control guidelines (*R*^2^ of linear regression on positive probe data below .95) and the other having a high positive correlation between *a*_*i*_ and *b*_*i*_ estimates, contrary to the rest of the pool. This left us with *K* = 13 datasets, which ranges in size from *n*_13_ = 8 up to *n*_9_ = 1, 950 samples. All 13 datasets had 6 positive probes, each having the same control RNA levels (128, 32, 8, 2, .5, .125 fM for probes 1-6, respectively).

### 2.2. Design

This section outlines the hierarchical model used to create our meta-analysis prior. We assume that the log_10_ count of each sample/probe/study combination *Y*_*ijk*_ is characterized by

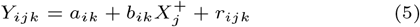

where 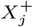 is the log_10_ RNA level for each probe *j*, which is fixed across all samples and studies at log_10_(128, 32, 8, 2, .5, .125), *a*_*ik*_ and *b*_*ik*_ are the slope and intercept terms for sample *i* of study *k*, and *r*_*ijk*_ is the residual term reflecting the remaining variability of *Y*_*ijk*_ after taking into account the linear trend, for *i* = 1, … *I*_*k*_, *j* = 1, … 6, and *k* = 1, … *K* (*I*_*k*_ is the number of samples in study *k* and *K* = 13). This structure maintains the original design of RCRnorm while accounting for multiple data sources.

First, we focus on modeling the random regression coefficients *a*_*ik*_ and *b*_*ik*_ in (5). Unlike RCRnorm, the meta-analysis includes samples from a variety of datasets, meaning heterogeneity between data sources is a potential factor. To examine this, we first calculated empirical estimates for all *a*_*ik*_, *b*_*ik*_ by fitting individual linear regressions of the form

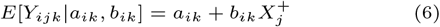

where each 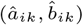 is calculated with 6 positive control observations of sample *i* (*j* = 1, …, 6). Figure 2(a) and (b) summarize the estimates obtained from this analysis. There is clear heterogeneity in the distributions, both in the center and variance, when broken down by dataset. Therefore it is reasonable to assume that parameters associated with *a*_*ik*_ and *b*_*ik*_ are diverse for different datasets. This leads us to a separate bivariate normal distribution in (7) for each study *k*, where the study-specific means *α*_*k*_ and *β*_*k*_ are equivalent to *μ*_*a*_, *μ*_*b*_ in the RCRnorm model.

The next step is to determine the nature of the covariance matrix Σ^(*k*)^, that is, whether or not *a*_*ik*_ is independent from *b*_*ik*_. Figure 2(c) shows a breakdown of correlation between the empirical estimates *â*_*ik*_ and 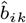 by dataset. Several datasets show positive correlation while others have a negative correlation, providing justification for modeling the study-specific correlation rather than assuming a common correlation value across all studies. Despite the fact that a few datasets would not have enough evidence to reject *r* = 0 at the *α* = .05 level, 12 of the 17 datasets have 95% confidence bounds strictly above or below 0, meaning we cannot assume that *a*_*ik*_ and *b*_*ik*_ are independent of each other when coming from the same dataset *k*. The implication of this is that Σ^(*k*)^ must be modeled with a multivariate distribution rather than independent univariate distributions. Several candidate distributions were considered, but a thorough simulation showed little difference in performance between prior distributions (interested readers can find results of this simulation in section S3 of the supplementary material [1]). We therefore chose the path of least resistance by implementing a classic inverse-Wishart distribution in (8), which is the conjugate prior for the covariance matrix of a multivariate normal distribution and allows for both simplicity and computational ease. Finally, the mean parameters *α*_*k*_ and *β*_*k*_ are modeled independently, since the empirical estimates have a correlation near 0. The hyperparameters *μ*_*α*_, *μ*_*β*_ have “safe range” uniform priors while 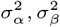 have diffuse inverse gamma priors. Safe range prior distributions are those that follow a uniform distribution with a range that comfortably covers all plausible values for the parameters (usually it has a half-width of 3 or 5 standard deviations).

The full hierarchical structure of the slope and intercept terms is summarized below,

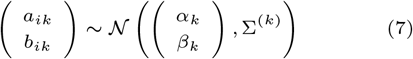

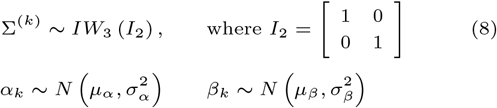

where *IW*_3_ is an inverse-Wishart distribution with three degrees of freedom.

We are now left with the residual term *r*_*ijk*_ from (5), which is modeled by the following hierarchical structure:

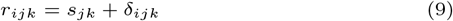

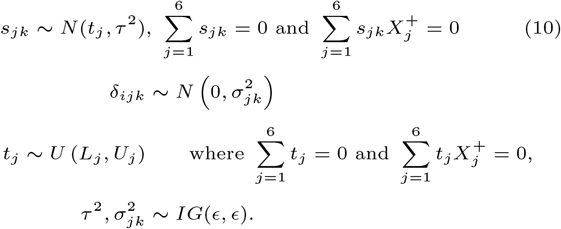

The above structure was carefully chosen to reflect what was observed from the data and to achieve computing efficiency and stability. Figure 2(d) shows the mean residual for each corresponding probe by dataset. While some probes seem to be centered at 0, there is a clear pattern in the residuals especially for 5 and 6. This indicates that the model for *r*_*ijk*_ should have probe specific terms. Since there is significant heterogeneity in the probe effect between different datasets, the parameters are specific to both the probe and dataset (i.e., indexed by *j, k*). Therefore the residual term can be broken down into two pieces, shown in (9). The parameters *s*_*jk*_ are assumed to have a probe specific mean *t*_*j*_ and a shared variance *τ* ^2^, as the points by probe in Figure 2(d) have centers clearly distinct from one another but have similar variability. We further enforce the constraints shown in (10), to avoid issues with identifiability, making *s*_*jk*_ dependent on other probe effects from the same dataset. We include these constraints to stabilize our distribution in the absence of strong prior information, relying on the fact that the log-transformed positive probe data have a strong linear relationship with the log-transformed control RNA expression levels in each study alone. Since these terms represent probe specific residuals, we can reasonably expect them to follow the same rules as residuals, namely that they sum to 0 with and without multiplication by the factor-effects. While this is a frequentist technique, there is significant value in using both frequentist and bayesian approaches together, despite the tendency to think of them as separate, binary categories [2]. In this case, our analysis benefits from the constraints by reaching quick convergence without using artificially informative prior distributions. A “safe range” uniform prior is applied to *t*_*j*_ (including the same constraints used for *s*_*jk*_) while a non-informative *IG* prior is used for *τ* ^2^. Finally, *δ*_*ijk*_ acts as our pure residual term that has no pattern remaining, which we center at 0. Given the diversity shown in this section by probe and by dataset, we assume that the variance of this residual differs by these factors. We use a diffuse *IG* prior for the 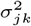 terms.

**Fig. 1.**
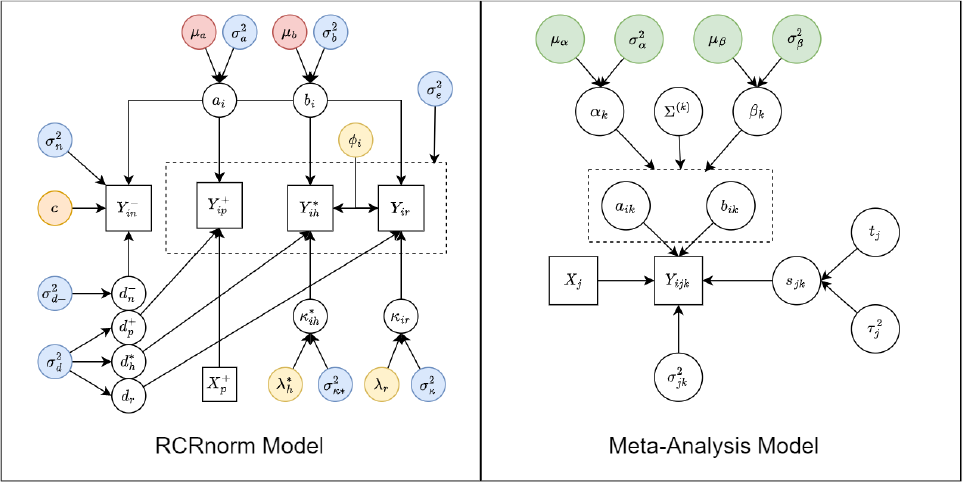
Graphic representation of the original RCRnorm model (left) and the meta-analysis model (right). Squares denote observed or fixed values while circles represent model parameters (random variables). Parameters and hyperparameters with blue color-coding represent diffuse inverse gamma priors, orange and yellow have “safe range” uniform priors, with yellow having ranges determined by positive probe data. Red hyperparameters have priors created by a jackknife analysis of the positive probe data. On the right, green parameters represent those being estimated by the meta-analysis.

**Fig. 2.**
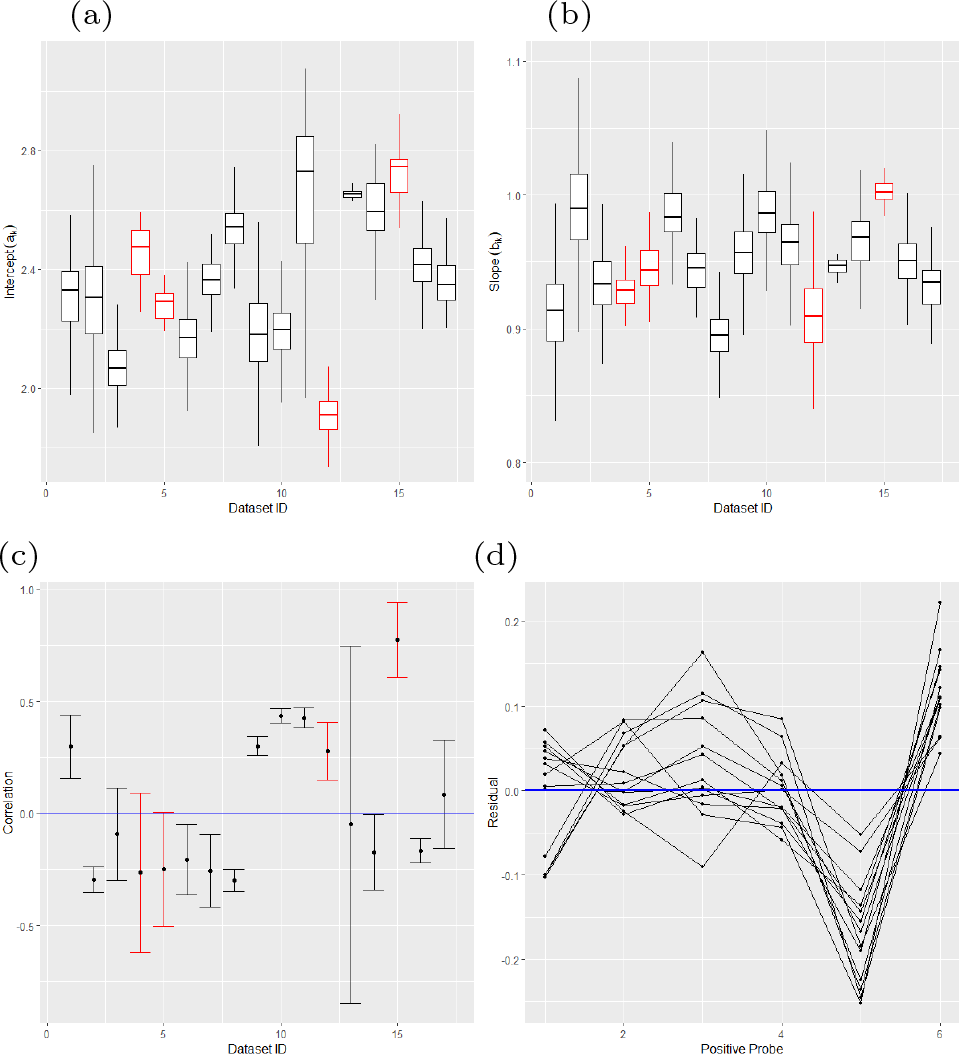
Meta-analysis: (a) and (b) show boxplots without outliers for empirical *a*_*ik*_ (left) and *b*_*ik*_ (right) estimates by dataset; (c) shows error bar plots of (*a*_*ik*_, *b*_*ik*_) correlation by dataset; (d) shows average empirical residual for positive probes by dataset. Red color represents datasets that were excluded from the meta-analysis (id 4, 5, 12, and 15). In plot (c), the dots represent the Pearson correlation between empirical estimates of *a*_*ik*_, *b*_*ik*_ while the bars reflect 95% confidence bounds. In plot (d), points on the same line come from the same dataset, and this plot only shows the datasets included in the meta-analysis.

The right panel of figure 1 gives a concise summary of variable relationships in the meta-analysis. The values we want to identify are represented by the green parameters in the figure. The full probability model and derivation of the full conditional distributions can be found in Section S1 of the supplementary material [1].

### 2.3. Diagnostics and Results

The MCMC algorithm for posterior sampling was written in R (R core team, 2019) and implemented via a Gibbs sampler using the package rcpp [6], which greatly reduced the computational cost. The meta-analysis model was run via 4 parallel chains of 12,000 samples, with each chain needing no more than 5,000 samples to converge (see figure S1 in the supplementary material [1] for trace plots for the global parameters).

For each parameter, the trace plots show that all four chains converge to the same distribution, which is further supported by the Gelman-Rubin diagnostics (median and 95% potential scale reduction factor at 1.00 for all four global parameters). Table 1 shows summary statistics for the posterior draws across all chains. Since we observed relatively low correlation among sequential draws, with all variables below 5% correlation for a lag of 2, we thin the chains after burn-in by 1 draw, so every other sample is used to calculate the summary statistics (a total of 14,000 samples). Therefore, we accept our meta analysis prior for *μ*_*a*_ and *μ*_*b*_ to be

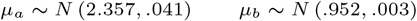

based on posterior means for *μ* parameters and posterior medians for *σ*^2^ parameters.

### 2.4. Additional Enhancements

Three major changes were made to the Gibbs sampler besides the informative prior. These first of these is implementing constraints on the *d*_*p*_ and *d*_*n*_ parameters to improve model convergence. While this may seem counterintuitive, such identifiability constraints have been shown to dramatically improve convergence of Gibbs samplers for Gaussian mixture models [19]. The other changes included updating *λ*_*r*_, *λ*_*h*_ and *ϕ*_*i*_ with fixed calculations and editing the underlying code to improve computational efficiency. A detailed description and justification of these changes can be found in section S4 of the supplementary material [1].

## 3. Application and Comparison to RCRnorm

We compare the performance and diagnostics of MetaNorm to that of RCRnorm in 5 different areas, including computation time, convergence, stability of estimates, model bias and performance with low-quality datasets. We will use four different datasets of varying quality, three that were excluded from the meta-analysis and one with a low number of samples. These datasets are summarized in Table 2. Dataset 4 is the lung cancer FFPE dataset used to evaluate RCRnorm [9] and contained in the RCRnorm R package. It contains 28 samples and 104 genes after cleaning, and is thought to be a high-quality dataset. Dataset 12 was excluded from the meta-analysis due to a high number of low-quality samples (low R-squared in the positive probe data) and dataset 15 was excluded for having a very high positive correlation between *a*_*i*_ and *b*_*i*_ (see figure 2(c)). Despite being included in the meta-analysis, Dataset 13 is also used for testing due to its low number of samples and high number of genes.

**Table 1.** Summary statistics for global parameters in our Bayesian meta-analysis after a burn-in of 5,000 and thinning every other draw.

| Parameter | Q1 | Median | Q3 | Mean | Std. Dev. |
| --- | --- | --- | --- | --- | --- |
| $\mu_\alpha$ | 2.318 | 2.357 | 2.395 | 2.357 | 0.060 |
| $\mu_\beta$ | 0.941 | 0.952 | 0.962 | 0.952 | 0.016 |
| $\sigma_\alpha^2$ | 0.031 | 0.041 | 0.056 | 0.047 | 0.024 |
| $\sigma_\beta^2$ | 0.002 | 0.003 | 0.004 | 0.003 | 0.002 |

**Table 2.**
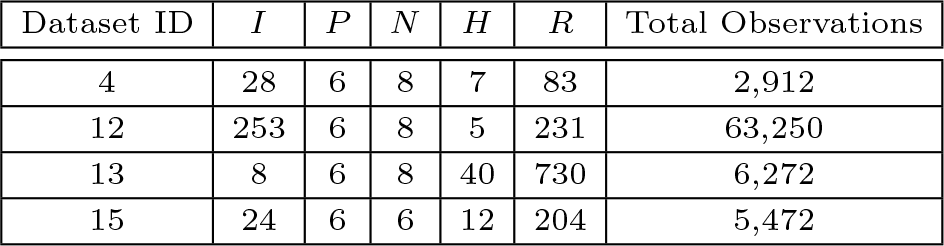
Characteristics of testing datasets.

| Dataset ID | $I$ | $P$ | $N$ | $H$ | $R$ | Total Observations |
| --- | --- | --- | --- | --- | --- | --- |
| 4 | 28 | 6 | 8 | 7 | 83 | 2,912 |
| 12 | 253 | 6 | 8 | 5 | 231 | 63,250 |
| 13 | 8 | 6 | 8 | 40 | 730 | 6,272 |
| 15 | 24 | 6 | 6 | 12 | 204 | 5,472 |

### Computation Time

After restructuring the algorithm, MetaNorm significantly improves the computation time. Table 3 shows a breakdown of computation time for different datasets and samples in R-studio using a laptop with an i7 core and 16 GB of RAM. The results show that MetaNorm outperforms RCRnorm in computational efficiency, with a minimum improvement of 6-fold. But the effects of MetaNorm is seen most clearly when the number of samples is large (*I >* 20), with RCRnorm taking 25 to 40 times longer to produce the same amount of samples. In addition to this, MetaNorm tends to reach convergence quickly with little autocorrelation, typically requiring *<*500 draws to reach a stationary distribution and leading to improvement of another 5-fold at least. So MetaNorm improves upon RCRnorm by being faster per-draw but also by requiring fewer draws, leading to an increase an efficiency by anywhere from 30-fold for low-sample datasets to 200-fold for large-sample datasets. Local memory storage also improves with MetaNorm, as shown in section 4.

**Table 3.** Comparison between RCRnorm and MetaNorm on computation time using four real datasets (1,000 draws per run)

| ID | Average Computation Time (seconds) |  | MetaNorm Improvement |
| --- | --- | --- | --- |
|  | RCRnorm | MetaNorm |  |
| 4 | 96.80 | 2.29 | 42.2 |
| 12 | 1340.60 | 39.39 | 34.03 |
| 13 | 55.86 | 9.26 | 6.03 |
| 15 | 90.88 | 1.78 | 25.60 |

**Table 4.** Comparison between RCRnorm and MetaNorm on (a) convergency using Gelman-Rubin diagnostics for *μ*_*a*_ and (b) median standard deviation of *κ*_*ir*_ s.

| | (a) Gelman-Rubin Diagnostics for $\mu_a$ | | | | (b) Median Standard Deviation of $\kappa_{i,r,s}$ | |
| --- | --- | --- | --- | --- | --- | --- |
| Dataset ID | RCRnorm |  | MetaNorm |  | RCRnorm | MetaNorm |
|  | Median | Upper C.I. (95%) | Median | Upper C.I. (95%) |  |  |
| 4 | 1.05 | 1.12 | 1.00 | 1.00 | 0.037 | 0.009 |
| 12 | 1.00 | 1.00 | 1.00 | 1.00 | 0.205 | 0.003 |
| 13 | 1.02 | 1.06 | 1.00 | 1.00 | 0.961 | 0.015 |
| 15 | 1.22 | 1.52 | 1.00 | 1.00 | 0.092 | 0.007 |

### Convergence

All datasets were run with 5 chains (randomized starting points) of 15,000 draws each except for dataset 12, which we limited to 5 chains of 5,000 draws each due to the size of the dataset. Convergence diagnostics for global parameter *μ*_*a*_ are shown in panel (a) of Table 4, where RCRnorm has decent convergence for dataset 4 and (to a lesser extent) dataset 15 but MetaNorm produces better convergence in either scenario. MetaNorm also converges in only a few hundred draws, much quicker than RCRnorm (see figure S5 [1]). Panel (a) of Table 4 also shows that datasets 12 and 13 achieve good convergence for global parameters regardless of which model is used. However the next level of the hierarchy tells a different story. Figure S6 in the supplementary material [1] shows trace plots of *a*_1_, *a*_2_, *a*_3_ from the first three patients for the RCRnorm (top) and MetaNorm (bottom) chains. Clearly, RCRnorm is not converging to the same distribution in every chain, which is a significant issue. MetaNorm resolves this issue, with all 8 samples converge in their *a*_*i*_ estimates. Consequentially, the normalized gene read counts vary significantly between runs with different starting points, casting doubt on the results (see panel (b) of Table 4). The issues raised in this section indicate that RCRnorm suffers from a kind of “faux” convergence, where good convergence around global parameters masks weak or absent convergence in lower-level parameters. It should be noted that this does not occur in all datasets when using RCRnorm - such as dataset 4 - but (to our knowledge) does not occur at all when using MetaNorm.

### Stability of κ_ir_ Estimates

The most critical piece of both the RCRnorm and MetaNorm models are the *κ* estimates for housekeeping and regular genes. Representing the RNA amount after accounting for sample degradation, the *κ* parameters are the true model output and any value that the models offer lies in these estimates. With this in mind, if we run separate chains on the same dataset, ideally we will observe very similar estimates for these parameters. We will once again show that MetaNorm improves upon RCRnorm in this way, providing more stable estimates for these parameters. Panel (b) of Table 4 shows the median standard deviation between chains among all *κ*_*ir*_ estimates. For datasets 12, 13 and 15, which do not have all chains of sample-specific parameters converging to the same distribution, the stark difference in standard deviation should not be surprising; if the *a*_*i*_ and *b*_*i*_ terms are different, of course the *κ*_*ir*_ terms would be wildly different. But even when the data is good-quality and convergence is somewhat stable (as with dataset 4), there is still significant reduction in standard error in MetaNorm. The left panel of Figure 3 shows the densities of these standard errors.

### Bias of κ_ir_ Estimates

A small simulation study was performed to compare model bias of the *κ*_*ir*_ estimates between RCRnorm and MetaNorm. Data for the simulation was generated based on dataset 4 (lung cancer FFPE), under constraints 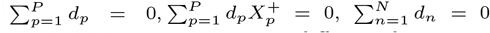. Bias was estimated using average mean difference between actual *κ*_*ir*_ values that the data was generated from and *κ*_*ir*_ estimates from the normalization procedures. In 100 runs of RCRnorm, we found that the normalization produced output that was off by an average of -.0651 per *κ*_*ir*_. This translates to an average 13.9% reduction from actual to estimated normalized gene read counts. In 100 runs of MetaNorm, we found that the output differed by an average of .0018 per *κ*_*ir*_. This translates to an average increase of 0.4% from actual to estimated normalized gene read counts. While this does show that MetaNorm has superior bias to RCRnorm, this result is not as significant as the reduction in prediction variance (especially as it pertains to mean-squared error contribution).

### Performance with Messy Data

This section presents a specific example of MetaNorm producing more intuitive and interpretable estimates for data with a large number of low quality samples. Dataset 12 contains 61 (24%) samples which have an *R*^2^ *<* .95 from a linear regression of positive probe data. Since the nCounter system is designed to maintain very strong linear relationship between the log-transformed counts and log-transformed control RNA expression levels, NanoString’s recommendation is to discard any samples with *R*^2^ *<* .95. In this section, we included all 253 samples for testing. Three chains of 10,000 draws were run on each model, with the first 5,000 discarded as burn-in. The estimates for *b*_*i*_ (the sample-specific slope terms) were drastically different, as the right panel of Figure 3 shows, despite the fact that the jackknife prior implemented in RCRnorm keeps the *μ*_*b*_ samples from deviating off the empirical estimate. While the MetaNorm estimates are within a reasonable neighborhood of *μ*_*b*_, the RCRnorm estimates are much lower, with a significant portion of them (∼34%) less than 0 (implying a negative relationship between read counts and mRNA). As a point of comparison, empirical estimates for *b*_*i*_ from dataset 12 have a minimum of .829. While some variation from the empirical estimates is expected, the amount seen from the RCRnorm is counter-intuitive both to the original data and to the nCounter system in general. Since *b*_*i*_ drives the relationship between output and mRNA for sample *i*, this causes significant downstream effects by producing inaccurate *κ*_*ir*_ which will ultimately lead to biased analyses. Alternatively, MetaNorm is able to overcome the pitfalls of messy data and produces reasonably intuitive estimates for *b*_*i*_. For this reason, the *κ*_*ir*_ estimates from MetaNorm are more plausible than those from RCRnorm for this dataset.

**Fig. 3.**
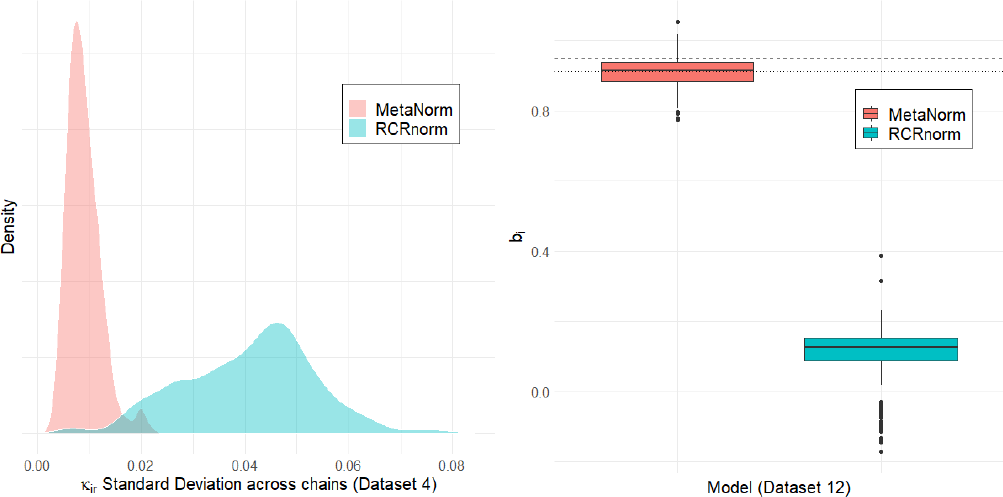
Comparison between RCRnorm and MetaNorm: the left panel shows the density of *κ*_*ir*_ ‘s standard errors for dataset 4; the right panel shows boxplots of *b*_*i*_ estimates for dataset 12, where the dashed line represents *μ*_*β*_, the meta-analysis estimate for the prior mean of *μ*_*b*_, and the dotted line represents the empirical estimate of *μ*_*b*_, using the positive probe data.

## 4. to RUVSeq

We conclude with a comparison to a leading normalization approach called RUVSeq [15], which is designed to remove unwanted technical variation. A recent study [3] compared the performance of RUVSeq with RCRnorm and two other approaches (nanostringdiff [18] and nSolver [12]) on their ability to remove technical variation and retain relevant biological variation from the Carolina Breast Cancer Study [13] (CBCS) dataset. While RCRnorm and RUVSeq outperform the other observed approaches, the computational load is much heavier for RCRnorm, requiring 20.93GB of local memory to obtain normalized estimates due to the size of the dataset (more than 500,000 total observations). Using the same dataset, we compare MetaNorm to RUVSeq on similar criteria.

The MetaNorm package contains an alternate function that runs the same normalization procedure, but keeps only the most recent draws for the *κ* parameters and calculates a rolling average for a posterior estimates. This allows researchers to avoid storing all draws for all parameters, which can cause major computational problems when large datasets are involved. Since the algorithm is continually freeing space by dumping old draws, the maximum amount of local memory needed is less than 1 GB for CBCS, making it much more comparable to approaches such as RUVSeq.

Each sample in the CBCS dataset is categorized by study phase (time period in which the sample was taken) which accounts for a significant amount of variability. Out of 406 regular genes included, only 6 (1.5%) did not show evidence for different means by study phase at a statistically significant level using an F-test on log-transformed read counts. When normalized estimates were used, these numbers jumped to 34 and 37 for MetaNorm and RUVSeq, respectively, suggesting that MetaNorm maintains good performance in the removal of technical variation as compared to RUVSeq. A similar analysis on gene expression variation due to ER (Estrogen-Receptor) status, finds that across raw data, MetaNorm and RUVSeq estimates, 77%, 90% and 86% of genes had statistically significant F-statistics. These results show that MetaNorm outperforms RUVSeq in retaining biologically relevant variation.

## 5. Discussion

When conducting statistical analysis in the Bayesian framework, many researchers rely on non-informative or diffuse prior distributions to maintain objectivity. However, when meaningful prior information exists and can be identified, using an informative prior distribution to accurately reflect current knowledge may lead to superior outcomes and great efficiency. This paper demonstrates a practical example of achieving improvement by incorporating such prior information via meta-analysis into data normalization.

RCRnorm is a groundbreaking normalization tool that improved upon existing methodology by leveraging the complexities of nCounter data. While other methods rely on frequentist techniques, the Bayesian framework allows RCRnorm to thoroughly explore the parameter space, identifying hidden information in the data, casting aside implausible assumptions, and increasing model transparency and interpretability. But despite its improvements, RCRnorm struggles to provide reliable estimates when the quality of data is in question. One reason for this is the “non-informative” data-based prior implemented for two of the most important global parameters *μ*_*a*_ and *μ*_*b*_, potentially reinforcing bias stemming from the data. The goal of this study was to enhance the RCRnorm model by (1) identifying prior distributions for *μ*_*a*_ and *μ*_*b*_ based on a comprehensive meta-analysis of FFPE datasets and (2) implementing additional algorithmic enhancements to improve computational cost, model convergence and stability of estimates. This new Gibbs sampler is called MetaNorm.

All datasets tested on MetaNorm for this manuscript converged in less than 1,000 draws, suggesting that smaller chains can be used for MetaNorm. We also showed that MetaNorm leads to more consistent convergence in global parameters while ensuring that sample-specific parameters (*a*_*i*_, *b*_*i*_) reliably converge to the same distribution from diverse starting points. MetaNorm significantly reduces the variance in normalized *κ*_*ir*_ estimates from those produced by RCRnorm, mitigating the risk of downstream issues beyond normalization. Finally, a dataset of triple negative breast cancer mRNA signatures shows that MetaNorm continues to produce realistic and intuitive parameter estimates when data-quality is a concern, while RCRnorm falters.

A significant downside to MCMC approaches in general is the amount of additional computation time needed to individually sample different parameters one after another. Estimating the posterior distribution with variational inference rather than MCMC would further improve computational efficiency and is a topic of future research [4].

## Footnotes

1 R Package: http://CRAN.R-project.org/package=NAPPA

