## Supplementary Material for "MetaNorm: Incorporating Meta-analytic Priors into Normalization of NanoString nCounter Data"

#### S1 Meta-Analysis Details

##### Full Probability Model

$$P(\boldsymbol{Y},\Theta) \propto \prod_{k=1}^{K}\left(\prod_{i=1}^{n_k}\left(\prod_{j=1}^{6} N\left(Y_{ijk}|a_{ik}+b_{ik}X_j^{+}+s_{jk},\sigma^2_{jk}\right)\right)\right)\cdot \prod_{k=1}^{K}\prod_{i=1}^{n_k}\left(\mathcal{N}\left(a_{ik},b_{ik}\middle|\begin{pmatrix}\alpha_k\\ \beta_k\end{pmatrix},\Sigma^{(k)}\right)\right)$$

$$\cdot\prod_{k=1}^{K}\left(N\left(\alpha_k\left|\mu_\alpha,\sigma^2_\alpha\right.\right)\cdot N\left(\beta_k\left|\mu_\beta,\sigma^2_\beta\right.\right)\cdot IW_3\left(\Sigma^{(k)}\left|I_2\right.\right)\right)\cdot\pi(\mu_\alpha)\cdot\pi(\mu_\beta)\cdot\pi(\sigma^2_\alpha)\cdot\pi(\sigma^2_\beta)$$

$$\cdot\prod_{k=1}^{K}\prod_{j=1}^{6}\left(N\left(s_{jk}\left|t_j,\tau^2\right.\right)\cdot\pi(\sigma^2_{jk})\right)\cdot\prod_{j=1}^{6}\left(\pi\left(t_j\right)\right)\cdot\pi(\tau^2)$$

##### Full Conditional Distributions

Let $\boldsymbol{\theta_{ik}}=\begin{pmatrix}a_{ik}\\ b_{ik}\end{pmatrix}$, $\boldsymbol{X_{ik}}=\begin{bmatrix}1 & X_{i1k}\\ ... & ...\\ 1 & X_{i6k}\end{bmatrix}$, $\boldsymbol{m_k}=\begin{pmatrix}\alpha_k\\ \beta_k\end{pmatrix}$, while $\boldsymbol{Y_{ik}},\boldsymbol{s_k}$ are vectors over all 6 $Y_{ijk}, s_{jk}$ and diag$(\sigma^2_{jk})$ is a diagonal matrix with $\sigma^2_{1k}...\sigma^2_{6k}$ in the diagonal. $I_{(L,U)}$ is the indicator function, where $I_{(L,U)}=1$ when $L<x<U$ and 0 otherwise. For all Inverse-Gamma (IG) distributions, $\epsilon=.01$.

$$\boldsymbol{\theta_{ik}}|\cdots,\boldsymbol{Y}\sim N\left(\mu_{\alpha\beta},\boldsymbol{\Sigma}^{(k)}_{\alpha\beta}\right)\text{ where}$$

$$\mu_{\alpha\beta}=\boldsymbol{\Sigma}^{(k)}_{\alpha\beta}\left(\boldsymbol{\Sigma}^{(k)^{-1}}\boldsymbol{m_k}+\boldsymbol{X^T_{ik}}\text{diag}(\sigma^2_{jk})^{-1}(\boldsymbol{Y_{ik}}-\boldsymbol{s_k})\right)\text{ and}$$

$$\boldsymbol{\Sigma}^{(k)}_{\alpha\beta}=\left(\boldsymbol{\Sigma}^{(k)^{-1}}+\boldsymbol{X^T_{ik}}\text{diag}(\sigma^2_{jk})^{-1}\boldsymbol{X_{ik}}\right)^{-1}$$

$$\boldsymbol{\Sigma}^{(k)}\Big|\cdots,\boldsymbol{Y}\sim IW_{n_k+3}\left(I+\sum_{i=1}^{n_k}\begin{bmatrix}(a_{ik}-\alpha_k)^2 & (a_{ik}-\alpha_k)(b_{ik}-\beta_k)\\ (a_{ik}-\alpha_k)(b_{ik}-\beta_k) & (b_{ik}-\beta_k)^2\end{bmatrix}\right)$$

$$\boldsymbol{m}_k|\cdots,\boldsymbol{Y}\sim BVN\left(\left(\boldsymbol{\Sigma}_m^{-1}+n_k\boldsymbol{\Sigma}^{(k)^{-1}}\right)^{-1}\left(\boldsymbol{\Sigma}_m^{-1}\boldsymbol{\mu}+\sum_{i=1}^{n_k}\left(\boldsymbol{\Sigma}^{(k)^{-1}}\boldsymbol{\theta}_{ik}\right)\right),\left(\boldsymbol{\Sigma}_m^{-1}+n_k\boldsymbol{\Sigma}^{(k)^{-1}}\right)^{-1}\right)$$

$$\mu_\alpha|\cdots,\boldsymbol{Y})\sim I_{(L_\alpha,U_\alpha)}\cdot N\left(\frac{1}{K}\sum_{k=1}^{K}\alpha_k,\frac{\sigma^2_\alpha}{K}\right)$$

$$\mu_\beta|\cdots,\boldsymbol{Y})\sim I_{(L_\beta,U_\beta)}\cdot N\left(\frac{1}{K}\sum_{k=1}^{K}\beta_k,\frac{\sigma^2_\beta}{K}\right)$$

$$\sigma^2_\alpha|\cdots,\boldsymbol{Y}\sim IG\left(\epsilon+\frac{K}{2},\epsilon+\frac{1}{2}\sum_{k=1}^{K}(\alpha_k-\mu_\alpha)^2\right)$$

$$\sigma^2_\beta|\cdots,\boldsymbol{Y}\sim IG\left(\epsilon+\frac{K}{2},\epsilon+\frac{1}{2}\sum_{k=1}^{K}(\beta_k-\mu_\beta)^2\right)$$

For $s_{jk}$, we apply two constraints and update the parameter vectors $s_{5k}$ and $s_{6k}$ based on $s_{1k}, s_{2k}, s_{3k}, s_{4k}$. Let $h_j = \frac{X_6 - X_j}{X_5 - X_6}$, $r_{ijk} = Y_{ijk} - a_{ik} - b_{ik}X_j$, $S_{1:4,-j} = \left(\sum_{l=1}^{4} s_{lk}\right) - s_{jk}$, $(XS)_{1:4,-j} = \left(\sum_{l=1}^{4} X_l s_{lk}\right) - X_j s_{jk}$.

$$s_{jk}|\cdots,\boldsymbol{Y} \sim N\left(\mu_{jk}^{(s)}, \sigma_{jk}^{2(s)}\right) \text{ where } j = 1,2,3,4$$

$$\sigma_{jk}^{2(s)} = \left(\frac{1}{\tau^2}\left(1 + h_j^2 + (1+h_j)^2\right) + \frac{n_k}{\sigma_{jk}^2} + \frac{n_k}{\sigma_{5k}^2}h_j^2 + \frac{n_k}{\sigma_{6k}^2}(1+h_j)^2\right)^{-1}$$

$$\begin{aligned}\mu_{jk}^{(s)} = \sigma_{jk}^{2(s)} &\left(\frac{t_j}{\tau^2} + \frac{(t_5 - \frac{X_6 S_{1:4,-j} - (XS)_{1:4,-j}}{X_5 - X_6})}{\tau^2}h_j - \frac{t_6 + S_{1:4,-j} + \frac{X_6 S_{1:4,-j} - (XS)_{1:4,-j}}{X_5 - X_6}}{\tau_6^2}(1+h_j)\right) \\ &+ \sigma_{jk}^{2(s)}\left(+\frac{1}{\sigma_{jk}^2}\sum_{i=1}^{n_k} r_{ijk} + \frac{1}{\sigma_{5k}^2}\sum_{i=1}^{n_k}\left(r_{i5k} - \frac{X_6 S_{1:4,-j} - (XS)_{1:4,-j}}{X_5 - X_6}\right)h_j\right) \\ &- \sigma_{jk}^{2(s)}\left(\frac{1}{\sigma_{6k}^2}\sum_{i=1}^{n_k}\left(r_{i6k} + S_{1:4,-j} + \frac{X_6 S_{1:4,-j} - (XS)_{1:4,-j}}{X_5 - X_6}\right)(1+h_j)\right)\end{aligned}$$

$$s_{5k} = \frac{X_6 \sum_{j=1}^{4} s_{jk} - \sum_{j=1}^{4} X_j s_{jk}}{X_5 - X_6}, \qquad s_{6k} = -\left(\sum_{j=1}^{5} s_{jk}\right)$$

For $t_j$, we apply two constraints and update the parameters $t_5$ and $t_6$ based on $t_1, t_2, t_3, t_4$. Note that $T_{1:4,-j}$ and $(XT)_{1:4,-j}$ are defined similarly to $S_{1:4,-j}$ and $(XS)_{1:4,-j}$ above.

$$t_j|\cdots,\boldsymbol{Y} \sim I_{(L_t,U_t)} \cdot N\left(\mu_j^{(t)}, \sigma_j^{2(t)}\right) \text{ where } t = 1,2,3,4$$

$$\sigma_j^{2(t)} = \left(\frac{K}{\tau^2}\left(1 + h_j^2 + (1+h_j)^2\right)\right)^{-1}$$

$$\mu_j^{(t)} = \sigma_j^{2(t)}\left[\frac{1}{\tau^2}\left(\sum_{k=1}^{K} s_{jk} + h_j\sum_{k=1}^{K}\left(s_{5k} - \frac{X_6 T_{1:4,-j} - (XT)_{1:4,-j}}{X_5 - X_6}\right) - (1+h_j)\sum_{k=1}^{K}\left(s_{6k} + T_{1:4,-j} + \frac{X_6 T_{1:4,-j} - (XT)_{1:4,-j}}{X_5 - X_6}\right)\right)\right]$$

$$\tau^2\big|\cdots,\boldsymbol{Y} \sim IG\left(\epsilon + 3K, \epsilon + \frac{1}{2}\sum_{k=1}^{K}\sum_{j=1}^{6}(s_{jk} - t_j)^2\right)$$

$$\sigma_{jk}^2|\cdots,\boldsymbol{Y} \sim IG\left(\epsilon + \frac{n_k}{2}, \epsilon + \frac{1}{2}\sum_{i=1}^{n_k}(Y_{ijk} - (a_{ik} + b_{ik}X_{ijk} + s_{jk}))^2\right)$$

#### S2 Tables and Figures for MetaNorm

Table S1: Meta-Analysis Datasets

| ID | Description of samples | # of samples | Reference | GSE ID or Link |
|---|---|---|---|---|
| 1 | Lung adenocarcinoma (LUAD) patients | 162 | Molania et al. (2019) | N/A[1] |
| 2 | Inflammatory bowel disease patients | 989 | Molania et al. (2019) | GSE73094 |
| 3 | Colon cancer patients | 96 | Chen et al. (2016) | GSE62932 |
| 4 | Lung cancer patients | 28 | Jia et al. (2019) | N/A[2] |
| 5 | Colorectal cancer (CRC) patients | 54 | Jia et al. (2019) | GSE86561 |
| 6 | Early stage CRC patient tumors | 144 | Low et al. (2017) | GSE81983 |
| 7 | Early stage CRC patient tumors | 131 | Low et al. (2017) | GSE81985 |
| 8 | Breast tumor samples | 1321 | Liu et al. (2016) | GSE74821 |
| 9 | Patients stimulated with anti-CD3/CD28 | 1950 | Molania et al. (2019) | GSE60341 |
| 10 | Healthy individuals (stimulated and controls) | 2441 | Molania et al. (2019) | GSE53165 |
| 11 | Carolina breast cancer study (CBCS) tumors | 1278 | Patel et al. (2022) | GSE148418 |
| 12 | Survivors of triple negative breast cancer | 254 | Cascione et al. (2013) | GSE45498 |
| 13 | Merkel-cell carcinoma patients | 8 | Gravemeyer et al. (2021) | GSE159662 |
| 14 | T-cell lymphoma or dermititis (plus controls) | 128 | Nielsen et al. (2019) | GSE143382 |
| 15 | Metastatic melanoma patients | 24 | DeVito et al. (2021) | GSE165745 |
| 16 | Breast cancer patients | 1253 | Pu et al. (2019) | GSE147126 |
| 17 | Squamous cell carcinoma patients | 67 | Meehan et al. (2020) | GSE148944 |

[1] Data can be found at https://github.com/RMolania/NanostringNormalization/tree/master/Example%201_LungCancerStudy
[2] Data can be accessed in the RCRnorm R package

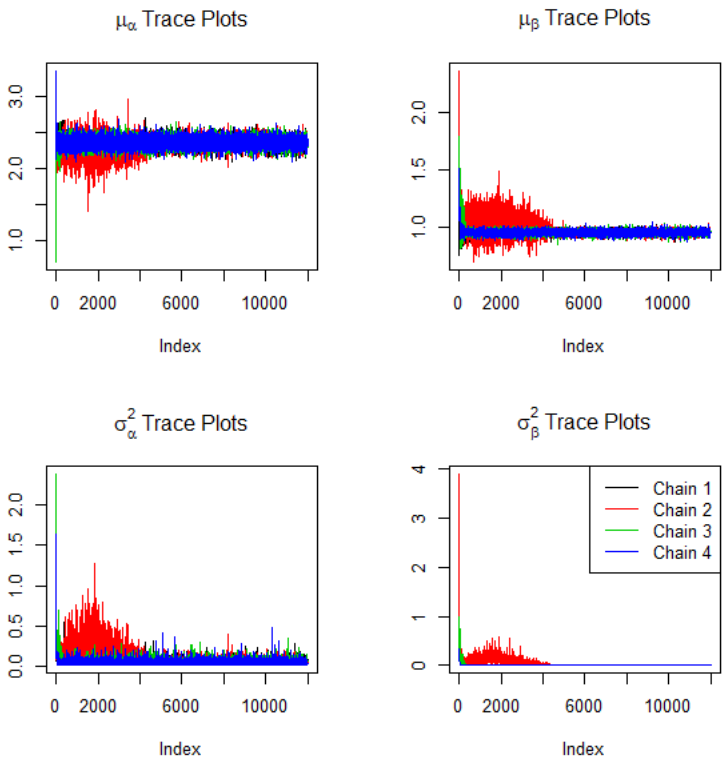


Figure S1: Trace plots (left) and Gelman-Rubin plots for global parameters in in our Bayesian meta-analysis

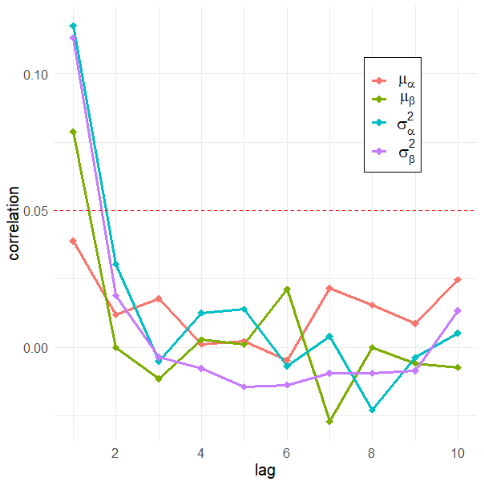


Figure S2: Autocorrelation for global parameters in our Bayesian meta-analysis. Since all chains produced similar results, only autocorrelation from chain 1 is shown.

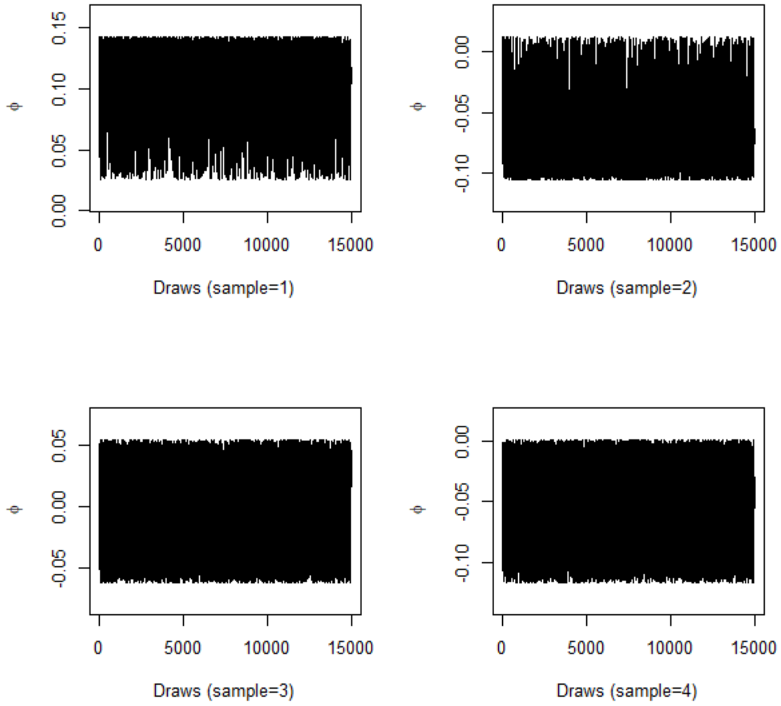


Figure S3: Trace plots for $\phi_1 - \phi_4$ (dataset 13).

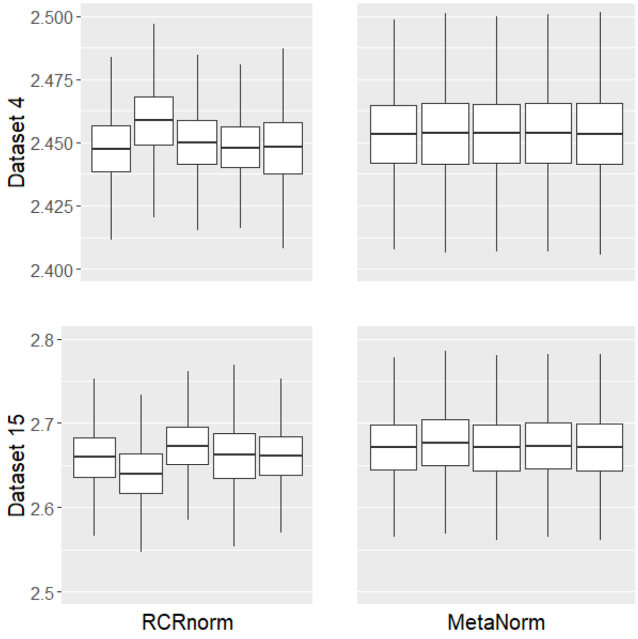


Figure S4: Posterior distribution (by chain) of $\mu_a$ for RCRnorm and MetaNorm normalization of datasets 4 and 15

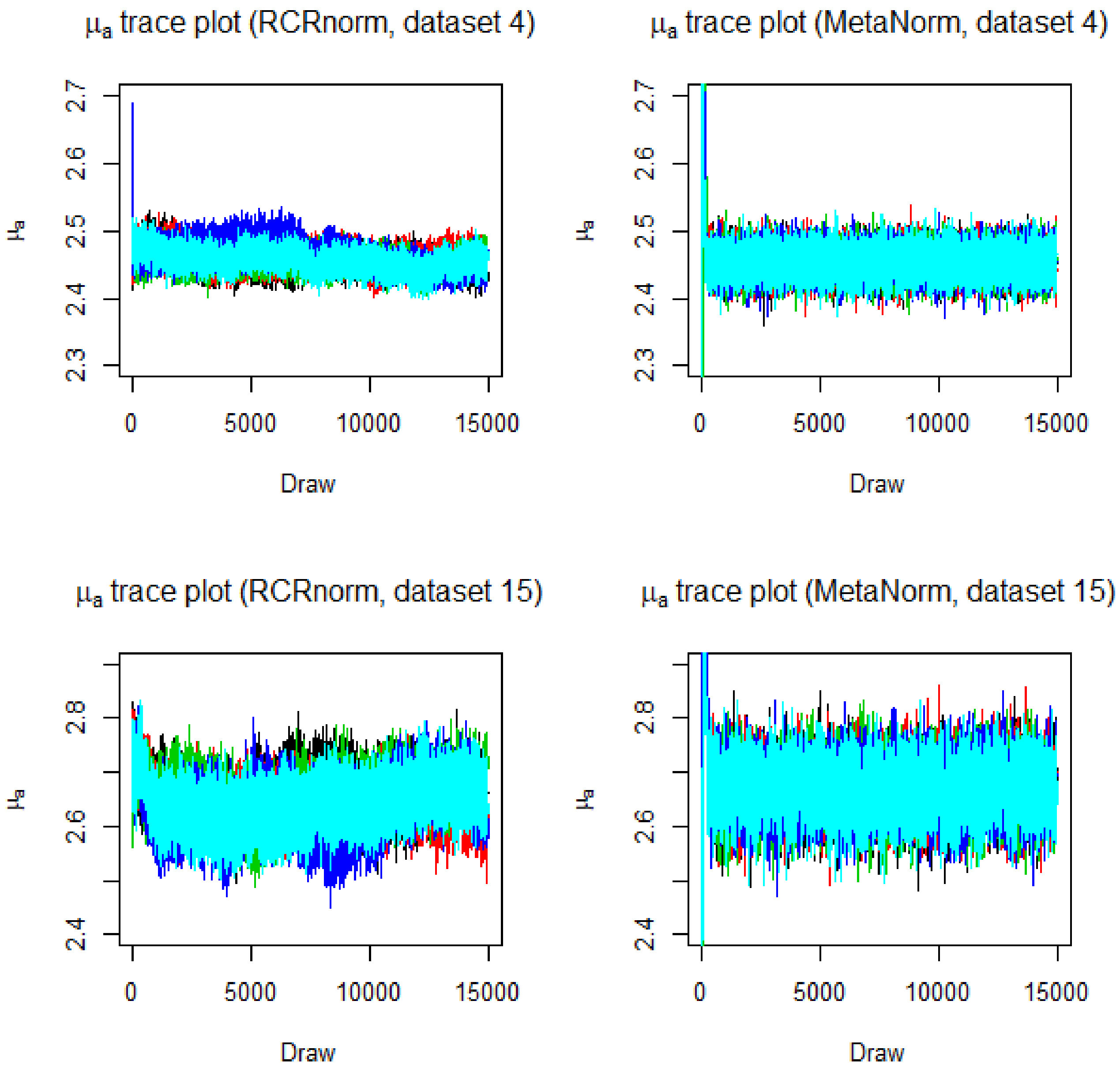


Figure S5: Comparison between RCRnorm and MetaNorm on convergency using trace plots for $\mu_a$

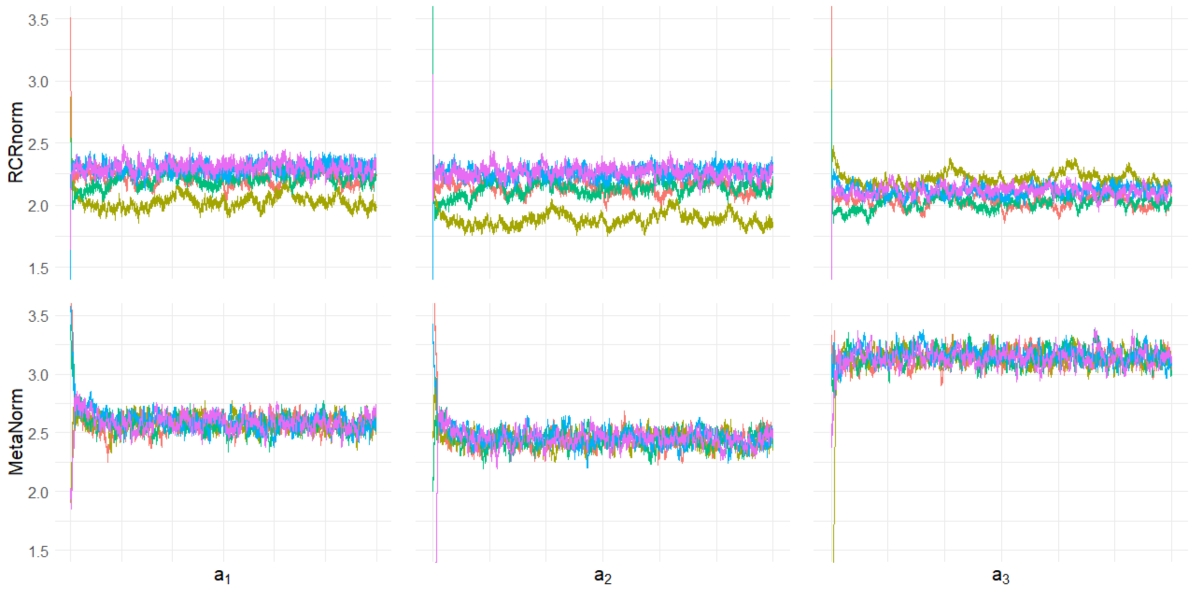


Figure S6: Comparison between RCRnorm and MetaNorm using traceplots of sample-specific intercepts $a_1 - a_3$ (Dataset 13).

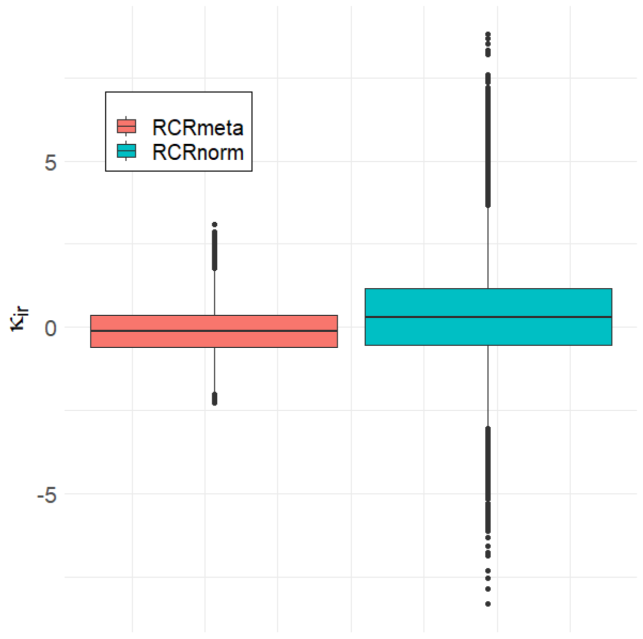


Figure S7: Comparison of $\kappa_{ir}$ (normalized $\log_{10}$ mRNA expression levels) for dataset 12

#### S3 Sensitivity Analysis

To confirm our choice of the prior distribution for the covariance matrix $\Sigma^{(k)}$, we conducted a simulation study and evaluated model performance in estimating the key global parameters $\mu_\alpha$ and $\mu_\beta$ across various objective priors for the bivariate covariance in the literature, as summarized in Berger and Sun (2008). Seven simulation settings were included in our study, each of which contains 50 synthetic datasets. For each dataset, we estimated $\mu_\alpha$ and $\mu_\beta$ using nine different covariance prior setups: Inverse-Wishart, Scaling, Half-t distribution, Prior J (Jeffery's prior), Prior IJ (Independence Jeffery's prior), Prior $R\rho$, Prior $R\sigma$, Prior $\hat{R}\sigma$, and Prior S. Details about each of these priors can be found in Berger and Sun (2008). Below we list the details of each simulation setting:

- Base: The empirical estimates of the model parameters are used as the truth.
- High Variance (HV): Parameters from the Base setting were used with the magnitudes of $\sigma_\alpha^2$ and $\sigma_\beta^2$ being multiplied by 2.
- Low Variance (LV): Parameters from the Base setting were used with the magnitudes of $\sigma_\alpha^2$ and $\sigma_\beta^2$ being divided by 2.
- Large Meta-Analysis (LMA, i.e., a large number of studies $K$): Parameters from the Base setting were used. However, the number of studies is doubled.
- Small Meta-Analysis (SMA, i.e., a small number of studies $K$): Parameters from the Base setting were used. However, the number of studies is halved.
- Large Sample (LS): Parameters from the Base setting were used. However, the number of patients within each study is doubled.
- Small Sample (SS): Parameters from the Base setting were used. However, the number of patients within each study is halved.

To add some more variation into our synthetic datasets, for each setting, we simulated the $\Sigma^{(k)}$'s. Specifically, the variance components of $\Sigma^{(k)}$'s were generated from Inverse-Gamma distributions whose parameters were estimated using moment matching. To simulate the correlation components, we first generated samples from a normal distribution with mean and variance based on observed correlations after Fisher's z-transformation.

The variables were then transformed back to the (-1,1) scale. Additionally, we generated the variance for each probe within each study from an Inverse-Gamma distribution where the parameters were estimated using moment matching. Finally, for each study, the number of samples ($n_i$) was drawn with replacement from the original numbers of patients.

For each covariance setup, we ran an MCMC chain of length 2,500 and burnt in the first half of the chain. To measure the estimation performance of each method, we calculated the MSEs of the estimated posterior means of $\mu_\alpha$ and $\mu_\beta$. The Scaling method took too long to generate posterior sample, so we excluded it from the comparison. The results, scaled by a factor 1000, are summarized in Table S2, which shows that in each simulation setting, the MSE's under the different priors are quite close to each other. Furthermore, there is no clear-cut winner among the priors. For instance, the inverse-Wishart approach has the highest MSE in the base scenario for $\mu_\alpha$ but has the lowest MSE in the same scenario for $\mu_\beta$. These results show that the meta-analysis performance has relatively low sensitivity to the prior choice for $\Sigma^{(k)}$.

| | Estimation of $\mu_\alpha$ | | | | | | | Estimation of $\mu_\beta$ | | | | | | |
|---|---|---|---|---|---|---|---|---|---|---|---|---|---|---|
| | Base | HV | LV | LMA | SMA | LS | SS | Base | HV | LV | LMA | SMA | LS | SS |
| IW | 2.522 | 4.392 | 1.259 | 1.162 | 7.941 | 3.09 | 4.072 | 7.741 | 11.08 | 4.206 | 3.347 | 12.11 | 6.288 | 7.489 |
| Half-t | 2.456 | 4.411 | 1.253 | 1.171 | 8.187 | 3.067 | 4.097 | 7.989 | 10.63 | 3.898 | 3.296 | 12.06 | 5.712 | 7.208 |
| Prior J | 2.423 | 4.534 | 1.289 | 1.172 | 8.141 | 3.053 | 4.019 | 7.758 | 10.29 | 3.989 | 3.355 | 12.27 | 5.663 | 7.222 |
| Prior IJ | 2.408 | 4.473 | 1.278 | 1.15 | 8.171 | 3.087 | 4.024 | 7.975 | 10.01 | 4.014 | 3.193 | 12.41 | 5.776 | 7.437 |
| Prior $R\rho$ | 2.483 | 4.482 | 1.316 | 1.177 | 8.077 | 3.039 | 4.071 | 7.984 | 10.36 | 3.884 | 3.257 | 12.69 | 5.791 | 6.982 |
| Prior $R\sigma$ | 2.403 | 4.44 | 1.281 | 1.154 | 8.085 | 3.058 | 4.039 | 7.876 | 10.27 | 4.013 | 3.268 | 12.15 | 5.983 | 7.489 |
| Prior $\hat{R}\sigma$ | 2.443 | 4.447 | 1.286 | 1.177 | 8.113 | 3.081 | 4.064 | 8.029 | 10.62 | 4.056 | 3.277 | 12.47 | 5.869 | 7.429 |
| Prior S | 2.396 | 4.498 | 1.307 | 1.175 | 8.148 | 3.068 | 4.075 | 7.872 | 10.26 | 4.022 | 3.224 | 11.98 | 5.865 | 7.3 |

Table S2: Analysis of sensitivity to various prior choices of the study-specific covariance matrix $\Sigma^k$: 1,000×MSE is reported to evaluate the performance of estimating $\mu_\alpha$ and $\mu_\beta$. LV stands for large variance, SV for small variance, LMA for large meta-analysis, SMA for small meta-analysis, LS for large sample, and SS for small sample.

### S4 Additional Enhancements

This section outlines the additional enhancements made to RCRnorm to improve model efficiency, convergence and stability. In addition to the meta-analysis prior, three changes were made to the model:

1. Implementing the constraints $\sum_{p=1}^{P} d_p = 0$, $\sum_{p=1}^{P} d_p X_p^+ = 0$, $\sum_{n=1}^{N} d_n = 0$ in the positive and negative probe equations
2. Updating $\lambda_r, \lambda_h$ and $\phi_i$ with fixed calculations (i.e., no random draws) in each iteration
3. Refurbishing R code with more efficient data structures and procedures to improve computational cost

One other notable update is that MetaNorm does not allow non-randomized starting points. The remainder of this section gives details and justification for the changes listed above.

#### Constraints on $d_p, d_n$

In the presence of a high number of local parameters, implementing reasonable constraints can help to stabilize a Gibbs sampler. This is the case with RCRnorm, where random intercepts and slopes $(a_i, b_i)$ are assumed for each sample along with probe-specific effects. In a frequentist analysis of the positive probes, two constraints would be required to achieve a unique solution. Without these constraints, there are an infinite number of solutions, reflecting the fact that predictive power could be distributed in different ways to the probe or sample effects. While this is less severe in an MCMC algorithm, where we rely on thousands of "solutions" (draws) to form a posterior distribution, convergence can sometimes suffer in such a highly complex system such as RCRnorm, especially when the sample size $I$ is not large. This effect can be seen in the trace plots of $d_6^+$ from the lung cancer FFPE dataset ($I = 28$) used for testing in Jia et al. (2019), shown in Figure S8. The lefthand plot shows a standard run of RCRnorm. While eventually all chains reach a similar range, it takes more than 10,000 draws for this to occur. It is also clear that there is a significant amount of autocorrelation and variability between chains, causing issues of replicability when chains are not long enough. The righthand plot shows 5 chains of RCRnorm with the constraints added. Clearly all chains are quickly converging to the same distribution, with minimal autocorrelation present. These constraints not only help to stabilize $d_p^+$, but also with other key model parameters including $\kappa_{ir}$.

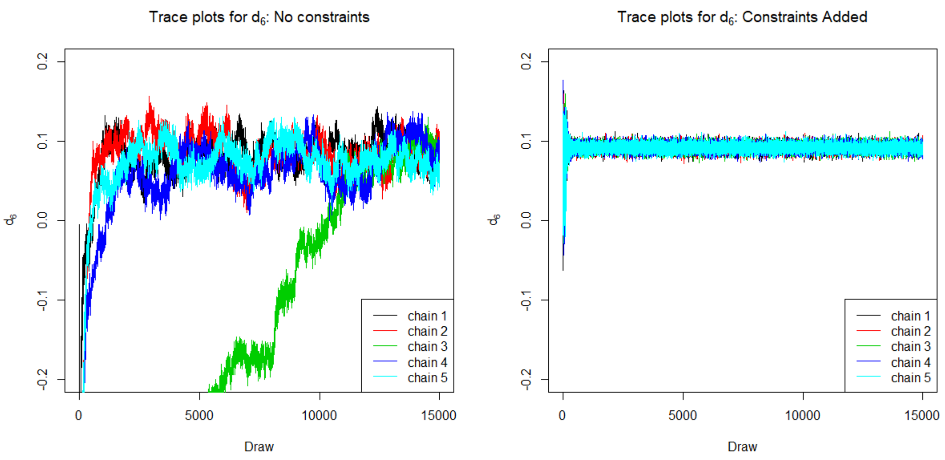


Figure S8: Trace plots of $d_6^+$ for RCRnorm with and without constraints (right and left, respectively) from the lung cancer FFPE dataset ($I = 28$) used for testing in Jia et al. (2019), showing that the convergence was greatly facilitated by the added constraints.

#### Updating $\lambda_h, \lambda_r$ and $\phi_i$

Now we turn our attention to the housekeeping and regular genes. Currently, RCRnorm relies on "safe range" uniform prior distribution for $\lambda_r, \lambda_h$ (the mean parameters for $\kappa_{ir}, \kappa_{ih}^*$ respectively) and $\phi_i$, which is generated based on empirical estimates of $a_i$ and $b_i$. Because there are a relatively large number of parameters for the information provided by the housekeeping and regular gene read counts, the samples for these parameters can oscillate within the pre-defined range, failing to converge to a much narrower distribution as we may expect. Figure S3 shows an example of this behavior using the $\phi_i$ trace plots from dataset 13 ($I = 8$).

Since the model structure is well-justified in Jia et al. (2019), we propose updating these parameters with a fixed calculation to stabilize the $\phi$ and $\lambda$ terms, while allowing the $\kappa$ terms to continue to be updated with its conditional distribution. First, we start by defining

$$\tilde{X}_{ih}^{(t)} = \frac{Y_{ih} - a_i^{(t)}}{b_i^{(t)}} \qquad \tilde{X}_{ir}^{(t)} = \frac{Y_{ir} - a_i^{(t)}}{b_i^{(t)}}$$

which is an estimate of the $\log_{10}$ RNA amount for housekeeping and regular genes given the $t^{th}$ draw of $a_i$, $b_i$. Let $j$ index both housekeeping and regular genes such that $j \in (1, ..., H, H+1, ..., H+R = J)$ so that $\tilde{X}_{ij}$ represents $\log_{10}$ RNA from both types of genes. For notational simplicity, we ignore the superscript $t$ here. We propose that the linear fixed-effects model $\tilde{X}_{ij} \sim \phi_i + \lambda_j$ with the constraint $\sum_{i=1}^{I} \phi_i = 0$ be estimated in every loop of the MCMC, with the parameter estimates $\hat{\phi}_i$ and $\hat{\lambda}_j$ used as the draws for $\phi_i, \lambda_j$. While this might seem computationally expensive, the factorial design of the data allows us to take a simple shortcut

$$\hat{\phi}_i = \frac{1}{J}\sum_{j=1}^{J} \tilde{X}_{ij} - \frac{1}{IJ}\sum_{i=1}^{I}\sum_{j=1}^{J} \tilde{X}_{ij} \qquad \hat{\lambda}_j = \frac{1}{I}\sum_{i=1}^{I} \tilde{X}_{ij}.$$

This method allows us to separate the effect of $a_i$ and $b_i$ (driven by the positive probes) so that $\phi_i$ can focus on the much more subtle sample effects in the housekeeping and regular gene counts. Similarly, this will allow us to obtain stable estimates of $\lambda_h, \lambda_r$ which will in-turn stabilize the $\kappa$ parameters. This will improve model convergence and allow the MetaNorm output to be significantly more precise (as shown in section 3 of Barth et al. (2023)), increasing its value and reliability to researchers.
